# Expanding the computable reactome in *Pseudomonas putida* reveals metabolic cycles providing robustness

**DOI:** 10.1101/139121

**Authors:** Juan Nogales, Steinn Gudmundsson, Estrella Duque, Juan Luis Ramos, Bernhard O. Palsson

**Affiliations:** Department of Bioengineering, University of California, San Diego, La Jolla, CA, USA; Department of Environmental Biology, Centro de Investigaciones Biológicas-CSIC, Madrid, Spain; Center for Systems Biology, University of Iceland, Iceland; Biotechnology Research Area, Abengoa Research, Sevilla, Spain

## Abstract

Genome-scale network reconstructions are organism-specific representations of metabolism and powerful tools for analyzing systemic metabolic properties. The use of reconstructions is limited by the lack of coverage of the metabolic reactome. We present an exhaustive and validated reconstruction of the biotechnologically relevant bacterium *Pseudomonas putida* KT2440, greatly expanding its computable metabolic states. The reconstruction, *i*JN1411, represents a significant expansion over other reconstructed bacterial metabolic networks. Computations based on the reconstruction exhibit high accuracy in predicting nutrient sources, growth rates, carbon flux distributions, and gene essentiality, thus providing a deep understanding of *Pseudomonas* metabolism. *i*JN1411 was used for: i) the assessment of the metabolic capabilities of *P. putida* as a species through multi-strain modeling, ii) deciphering the molecular mechanisms underlying metabolic robustness, and iii) identification of metabolic “capacitors” based on ATP-fueled metabolic cycles. This study represents the most complete and comprehensive bacterial metabolic reconstruction built to date, while providing computational and experimental evidence about how bacteria increase metabolic robustness, paving the way for engineering more robust biocatalysts and searching for drug targets in robust pathogens.

## Introduction

Robustness, understood as the property that allows systems to maintain their functions despite external and internal perturbations, is a systems-level phenomenon ubiquitously observed in nature, however it is still poorly understood at molecular level. There is therefore much interest in deciphering the role of biological robustness in research areas as diverse as multifactorial human diseases (1), evolution (2, 3), bacterial behavior (4), persistence (5), and biotechnology (6). Concepts from network theory, control theory, complexity science, and natural selection have been used to study robustness, however the molecular mechanism responsible for biological robustness remains uncertain (7). Four mechanisms ensuring biological robustness have been proposed (8). They included i) system control, e.g., the integral feedback between the components of a system allowing a coordinate response to perturbations; ii) alternative (redundant) mechanism, e.g., the presence of more than one identical, or similar, component which can replace the other, avoiding the collapse of the whole system when one of them fails; iii) network modularity, e.g., the spatial and temporal compartmentalization of biological networks, and iv) decoupling (non-specificity) e.g., the mechanism allowing biological networks to decouple low-level genetic variation from high-level functionalities.

Biological robustness in metabolic networks (metabolic robustness) has been studied in detail in yeast through gene-essentiality analysis (9) and flux analysis (10). Furthermore, <u>ge</u>nome-scale <u>n</u>etwork <u>re</u>constructions (GENREs) have been used to study the buffering capacity of metabolic networks against genetic perturbations (11, 12). More recently, the metabolic robustness under genetics and environmental perturbations has been addressed by using topological models highlighting the role of the network topology and the alternative (redundant) mechanisms underlying robustness (4). However, the systematic assignment of organism-specific biological robustness, at the genome-scale, will require the use of more complete and comprehensive computational models including key metabolic properties of the target organism extending beyond primary metabolism.

Through an exhaustive analysis of both the content and completeness of available GENREs we have recently revealed important shortcomings (13). We showed that i) a large space of the biological metabolic diversity across the phylogenetic tree has not yet been reconstructed, and ii) a significant portion of known species-specific reactions is missing from the corresponding reconstructions. As a result, most of current GENREs are highly similar in their reactomes content irrespective of their phylogenetic assignment, and thus they represent, roughly, models of primary metabolism rather than true organism-specific reconstructions. Compounding this limitation, the curation process of GENREs that involves resolving hundreds, even thousands, of ambiguities inherent to reaction properties, still lacks rigorous confidence standards and bibliomic support in most cases. Overall, these limitations hamper the use of <u>ge</u>nome-scale <u>m</u>odels (GEMs) in systems biology studies and an increasing number of researchers are calling for a more explicit curation process and the use of higher standards in the field (13–16). There are some notable exceptions that must be highlighted. The large metabolic reconstruction efforts on *E. coli* and yeast have resulted in the availability of high-quality reconstructions for these organisms (17, 18). The GEMs of *E. coli* have provided a better understanding of genotype-phenotype relationships in *E. coli* metabolism, (19, 20) while have unraveled systems properties of bacterial metabolism (21, 22). Therefore, similar efforts are required for other bacterial groups in order to expand the current biological reactome suitable for computation while providing a chance to unravel new bacterial systems properties such as metabolic robustness.

The group *Pseudomonas* comprises a heterogeneous and large group (> 100) of Gram-negative, gamma-proteobacterial species (23). They show a noteworthy metabolic versatility and adaptability enabling colonization of diverse niches (24). *Pseudomonas* are of great interest because of their importance in human and plant diseases, e.g., *P. aeruginosa* (25) and *P. syringe* (26), and due to their potential for promoting plant growth and biotechnological applications, e.g., *P. fluorescens* (27) and *P. putida* (28, 29). Among this group, *P. putida* has been widely used as a model environmental bacterium free of undesirable biotechnological traits such as virulence factors (30). *P. putida* strains can degrade a large array of chemicals, including xenobiotic compounds, while exhibiting a remarkable resistance to organic solvents and other environmental stresses which make *P. putida* strains highly-valued biocatalysts (31–34). In addition, *P. putida* strains are susceptible to genetic modification and are therefore seen by many as ideal workhorses for synthetic biology-based cell factories (34).

This high level of interest in *P. putida* has led to intense genome-scale metabolic modeling efforts of the strain KT2440; the best characterized strain and the first to be completely sequenced (35). Four GEMs for KT2440 have been previously published, formally known as *i*JN746 (36), *i*JP850 (37), PpuMBEL1071 (38), and *i*JP962 (39). These models have been used for studying metabolic features of *P. putida* such as Polyhydroxyalkanoate (PHA) and aromatic acids metabolisms. Recently, two new so-called consensus models, formally *i*EB1050 (40) and PpuQY1140 (41), have been published based on the genome reannotation of this strain and the integration of reactome already present in previous *P. putida* GEMs, respectively. Unfortunately, due the nature of this approach, which only allows the inclusion of new metabolic capabilities based on computational evidences with scarce experimental validation, and/or from previous reconstructions, the available GEMs of *P. putida* still lack coverage of the known metabolism in *P. putida* and fall into what we consider to be models of primary metabolism. Thus, as often occurs with current GEMs, their utility falls short of true and full genome-scale studies.

We show here that the entire metabolic knowledge available for a single species, even a genus, can be manually collected and used for high-quality metabolic modeling of a particular strain capable of addressing deep systems biology questions. We present a complete and manually curated metabolic reconstruction of *P. putida* KT2440, named *i*JN1411. This detailed reconstruction not only largely captures the metabolic features of this strain but it represents a computational scaffold for a future semi-automatic reconstruction of the *Pseudomonas* group. We use *i*JN1411 to develop a better understanding of metabolic robustness in bacteria, identifying processes non-essential for growth as responsible for this emergent property. Finally, we identify metabolic robustness cycles acting as metabolic capacitors responsible for connecting catabolism and anabolism with central metabolism.

## Results

### Reconstruction content and enhancements

The overall workflow for the reconstruction process is shown in SI1, (Fig. S1), and it is detailed in methods section. We followed a manual and iterative tri-dimensional approach based on i) genome annotation, ii) biochemical legacy knowledge, and iii) phenotypic experimental validation. As a result, a more accurate assignment of function to 246 genes was achieved (Table S1).

*i*JN1411, represents a significant expansion over previous GENREs from *P. putida* KT2440, and even over *E. coli* reconstructions (Table 1, SI1 Fig. S2). *i*JN1411 contains 1411 gene products (37% of the functionally annotated protein products in the KT2440 genome), 2826 reactions, and 2083 non-unique metabolites distributed over 90 specific subsystems over three different cellular compartments: extracellular, periplasm, and cytoplasm (Table S1). The reconstruction includes 409 unique citations and 2035 of the reactions have, at least, one citation supporting its inclusion (Table S1).

**Table 1.**
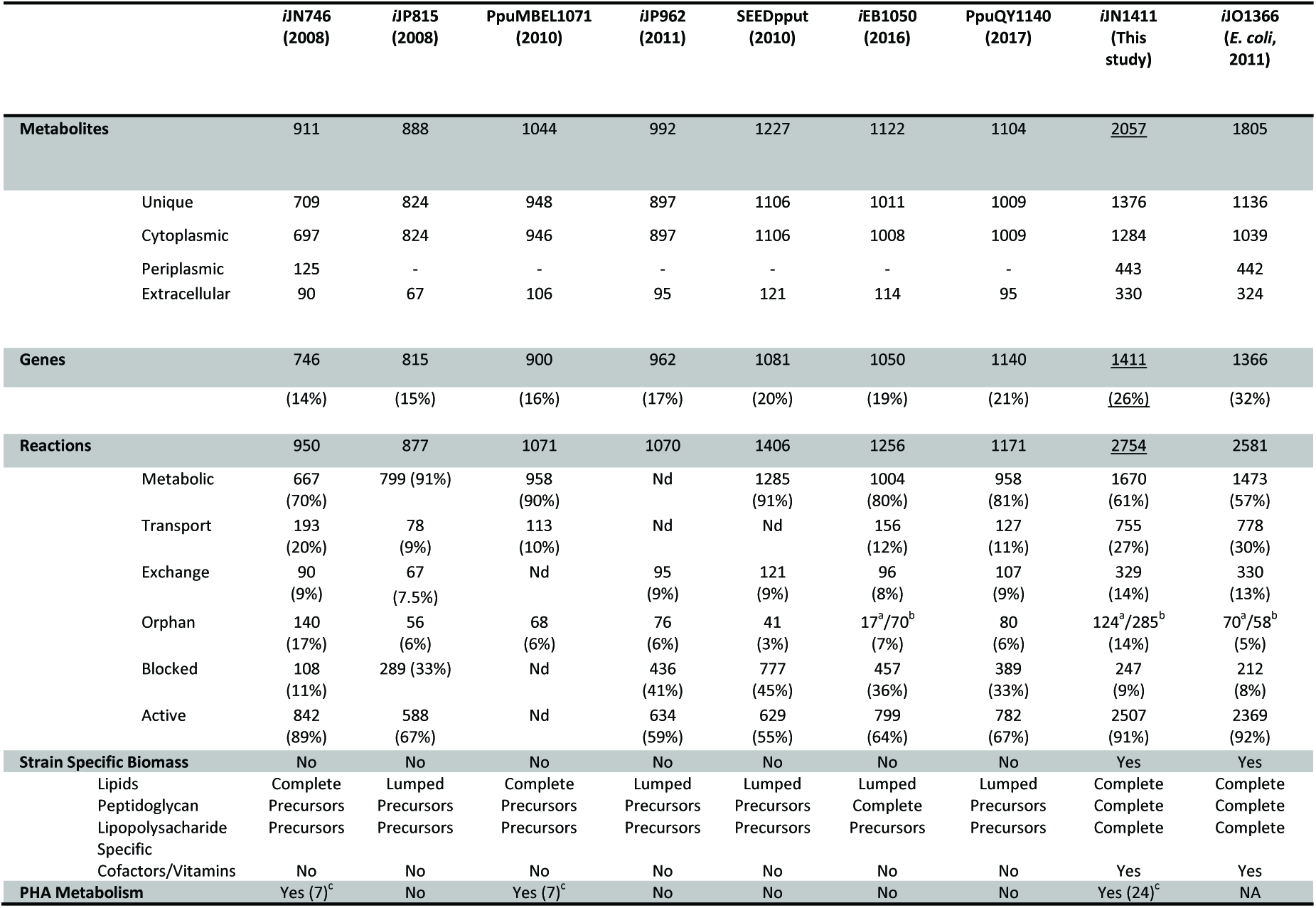
Comparison of the metabolic properties of *i*JN1411 with its antecessor *i*JN746 (36), previous *P. putida* metabolic reconstructions *i*JP815 (37), PpuMBEL1071 (38), *i*JP962 (95), the automatic reconstruction from SEED (87), and the recently published *P. putida* consensus models *i*EB1050 (40) and PpuQY1140 (41). Last *E. coli* reconstruction *i*JO1366 (17) was included as a reference for a high-quality GEM. ^a^Metabolic, ^b^Transport, ^c^Number of PHA monomers.

The major enhancements of *i*JN1411 over previous *P. putida* reconstructions are found in the strain-specific metabolism (Fig. 1A). New subsystems in *i*JN1411 account for well-known metabolic features of *P. putida*. For instance, stressors resistance is included in the subsystem heavy metal and solvent tolerance. The metabolic versatility of *P. putida* (42) has been captured in new subsystems such as alternate carbon and nitrogen sources. New catabolic pathways, many of which have been validated experimentally here, were included in *i*JN1411. For instance, the complete modeling of the sarcosine and 2,5 dioxopentanoate cataboloms, polyamines, and isovaleryl-CoA metabolisms have been included based on legacy data and completely validated by growth and gene knockout analysis (SI1, Fig. S3-6).

**Figure 1.**
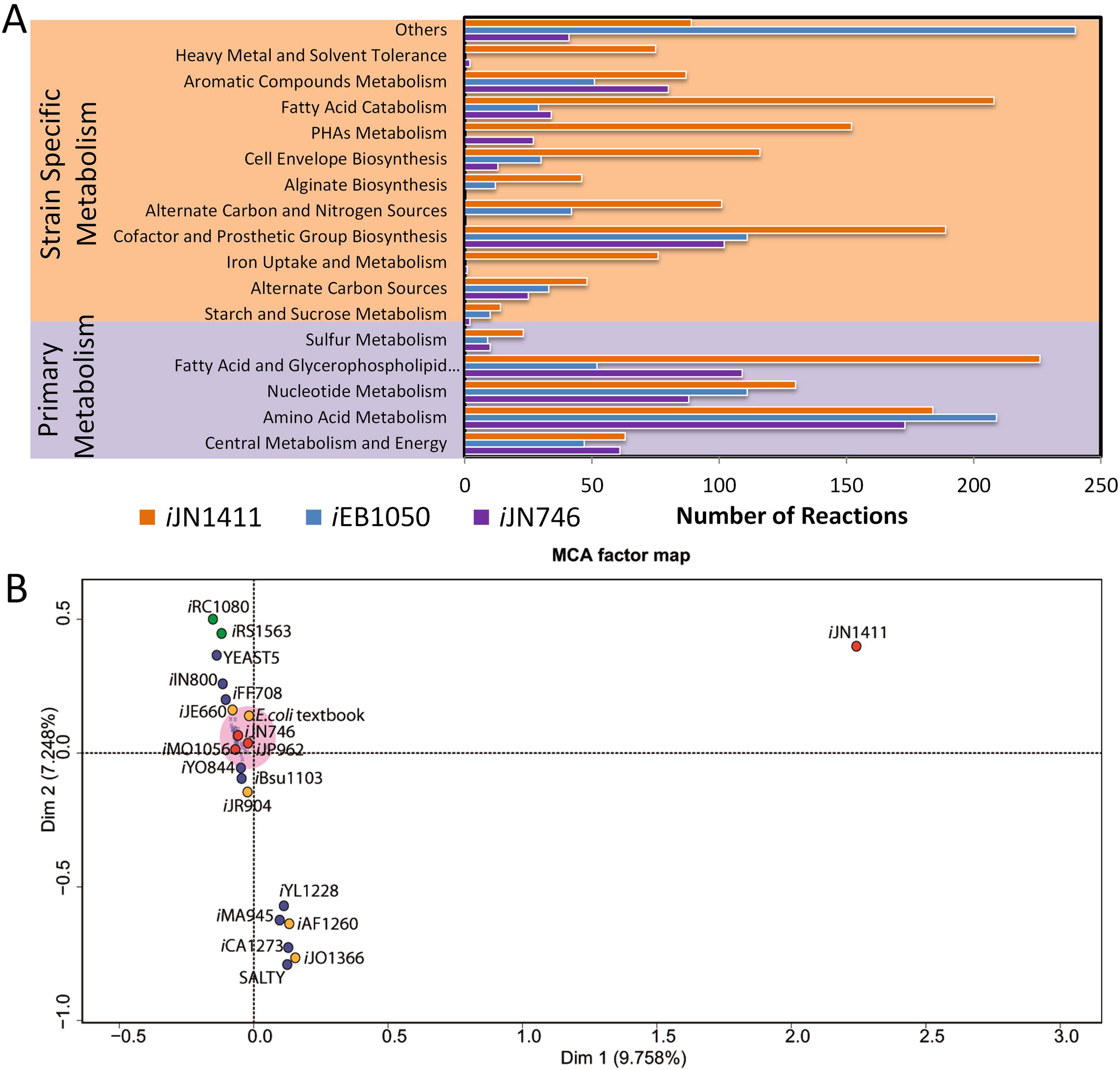
Metabolic content of *i*JN1411. **A. Metabolic content of *i*JN1411 categorized by subsystems compared with *i*JN746 and *i*EB1050.** Subsystems belonging to primary and strain-specific metabolisms are shaded in purple and orange, respectively. **B. Multiple correspondence analysis of the metabolic content, in terms of reactions and metabolites, of available GEMs (Monk et al, 2014) with *i*JN1411 added.** Models that are close to each other in the diagram are likely to have similar metabolic content. Most of the GEMs analyzed (shaded in pink), including *i*JN746 and *i*JP962 and iMO1056 (*P. aeruginosa* PA01), cluster around the origin, together with the reduced model of *E. coli* (*E. coli* textbook). Eukaryotic GEMs, including the model of *Chlamydomonas reinhardtii* (iRC1080), Zea mais (iRS1563) and yeast (YEAST5) are significantly different, including specific metabolic content. GEMs of *Enterobacteria* form a clearly differentiated group. Finally, *i*JN1411 is located far away from the origin, taking up a newly modeled metabolic space. Therefore, *i*JN1411 illustrates how different the metabolic space of *Pseudomonas* is compared to *Enterobacteria*. *P. putida*, *E. coli* MG1655, and other representative GEMs are indicated in red, orange, and blue, respectively. *i*IN800 and *i*FF708 (*S. cerevisiae*), *i*YO844 and *i*Bsu113 (*Bacillus subtilis*), *i*YL1228 (*Klebsiella pneumoniae*), *i*MA945 and SALTY (Salmonella Typhimurium), *i*CA1273 (*E. coli* W).

With regard to biosynthetic pathways, we performed detailed modeling of alginates, a *Pseudomonas* polysaccharide with high biotechnological and clinical interest. *Pseudomonas* has robust iron uptake metabolism that has a major role in niche colonization and pathogenesis (43, 44). Accordingly, the iron metabolism has been modeled including the biosynthetic pathway for pyoverdine (a non-ribosomal peptide acting as siderophore) of *P. putida* KT2440 based on structural studies (45).

The set of existing subsystems was significantly expanded (Fig. 1A). Within the cell envelope biosynthesis, specific peptidoglycans from *P. putida* and the complete lipopolysaccharide biosynthesis pathway have been modeled in great detail based on available data (46, 47). The modeling of the cellulose, rhamnose and trehalose metabolisms have been included as well. The biosynthesis for most of the cofactors and prosthetic groups known to be present in *Pseudomonas* was revisited in *i*JN1411, some of them, such as the biosynthesis of the pyrroloquinoline quinone (PQQ), are modeled here for the first time. These updates allowed for the assignment of the correct electron carrier to quinoproteins of *Pseudomonas* and a very accurate and strain-specific biomass reaction (see below).

*P. putida* catabolizes a large variety of fatty acids (48). Subsequently, the metabolism of fatty acids has been extensively expanded. In addition to saturated fatty acids, the catabolism of triacylglycerides, mono and poly-unsaturated fatty acids, phenylacyl, and thioacyl fatty acids, both with even- and odd-numbered chains, has been reconstructed. In addition, the metabolism of unsaturated fatty acids present in other bacterial models such as *i*JO1366 (17) has been revisited and extended by the inclusion of a NADPH-dependent 2,4-dienoyl-CoA reductase which is required for the β-oxidation of polyunsaturated fatty acids and substrate-specific *cis*-3-*trans*-2-enoyl-CoA isomerase reactions (48). As a direct consequence, the potential substrates for polyhydroxyalkanoate (PHA) synthesis via β-oxidation have experienced a significant increase and 24 different PHA monomers can be synthetized by *i*JN1411 (Fig. S7). Despite the production of PHA is one of the most prominent biotechnological capabilities of *P. putida*, the PHA metabolism is absent in most of the previous GEMs with the exception of *i*JN746 and PpuMBEL1071 (Table 1).

Finally, we proceed to construct a very detailed *P. putida*-specific biomass reaction (BOF) based on existing experimental data including macromolecular composition (49), glycerophospholipid content (50), murein composition (46), lipopolysaccharide (51), and specie-specific soluble metabolites such as pyoverdine (45) and pyroloquinolin quinone. A new value for non-associated growth maintenance (NGAM) was included as well based on recent findings (52). This highly strain-specific BOF contrasts with those present in previous reconstructions which lack *P. putida’s* specific lipids, lipopolysaccharides, peptidoglycans and the most of cofactors and vitamins (Fig. S8-9.) In addition, we formulated a core biomass reaction including those metabolites completely essential for growth according experimental reports. Details from the new biomass reactions and its formulation are depicted in SI1 and Table S3.

The metabolic expansion of the reactome represented by *i*JN1411 becomes evident when its content was compared with 53 GENREs by means of multiple correspondence analyses. While previous *P. putida* reconstructions such as *i*JN746 and *i*JP962 are located close to the center of coordinates together with most of the current GENREs, *i*JN1411 is placed far away from the center, illustrating its higher and organism-specific metabolic content (Fig. 1B). A single comparison with *i*JO1366 highlights this fact, showing that *i*JN1411 includes 1406 and 854 unique reactions and metabolites, respectively.

### Reconstruction validation through growth performance

To assess the ability of *i*JN1411 to predict physiological states, we first evaluated all the potential carbon, nitrogen, sulfur, phosphorus, and iron sources supporting *in silico* growth (Table S2). *i*JN1411 was able to use a significantly higher number of nutrients compared to previous reconstructions (Fig. 2). In fact, *i*JN1411 is able to grow on 140 and 71 new carbon and nitrogen sources, respectively, many of which never have been before experimentally reported as nutrients in *P. putida* (Table S2). Therefore, *i*JN1411 captures the metabolic versatility of *Pseudomonas* to a large extent. We then validated experimentally the accuracy of the growth predictions with special emphasis on those nutrients no tested so far in *P. putida* (SI1 Table S2). The overall accuracy of growth predictions was very high and it correctly predicted 80% and 82% (two-sided p-values of Fisher‘s exact test were less than 10^−13^) of the growth phenotypes observed for carbon and nitrogen sources, respectively (Fig. 2, Table S2). However, some discrepancies were found and they are discussed in detail in SI1. These discrepancies pave the way for reevaluating the metabolic versatility of *P. putida* in the context of silent metabolic pathways, underground metabolism, and/or unknown regulatory mechanisms. Comparisons of growth rate predictions and PHA production rates (Table 2) with experimental values provided further validation of the model. The prediction accuracy of *i*JN1411 significantly exceeds those from previous *P. putida* GEMs. However, *i*JN1411 grew faster than KT2440, suggesting an incomplete adaptation of KT2440 to these sugars as carbon sources and/or certain overflow of metabolism. In fact, when the observed secretion rates for gluconate and 2-ketogluconate were included in the model as additional constraints, *i*JN1411 fits the experimental growth rate on glucose. A similarly high level of accuracy was found for the growth rate and production rate of PHA on octanoate.

**Figure 2.**
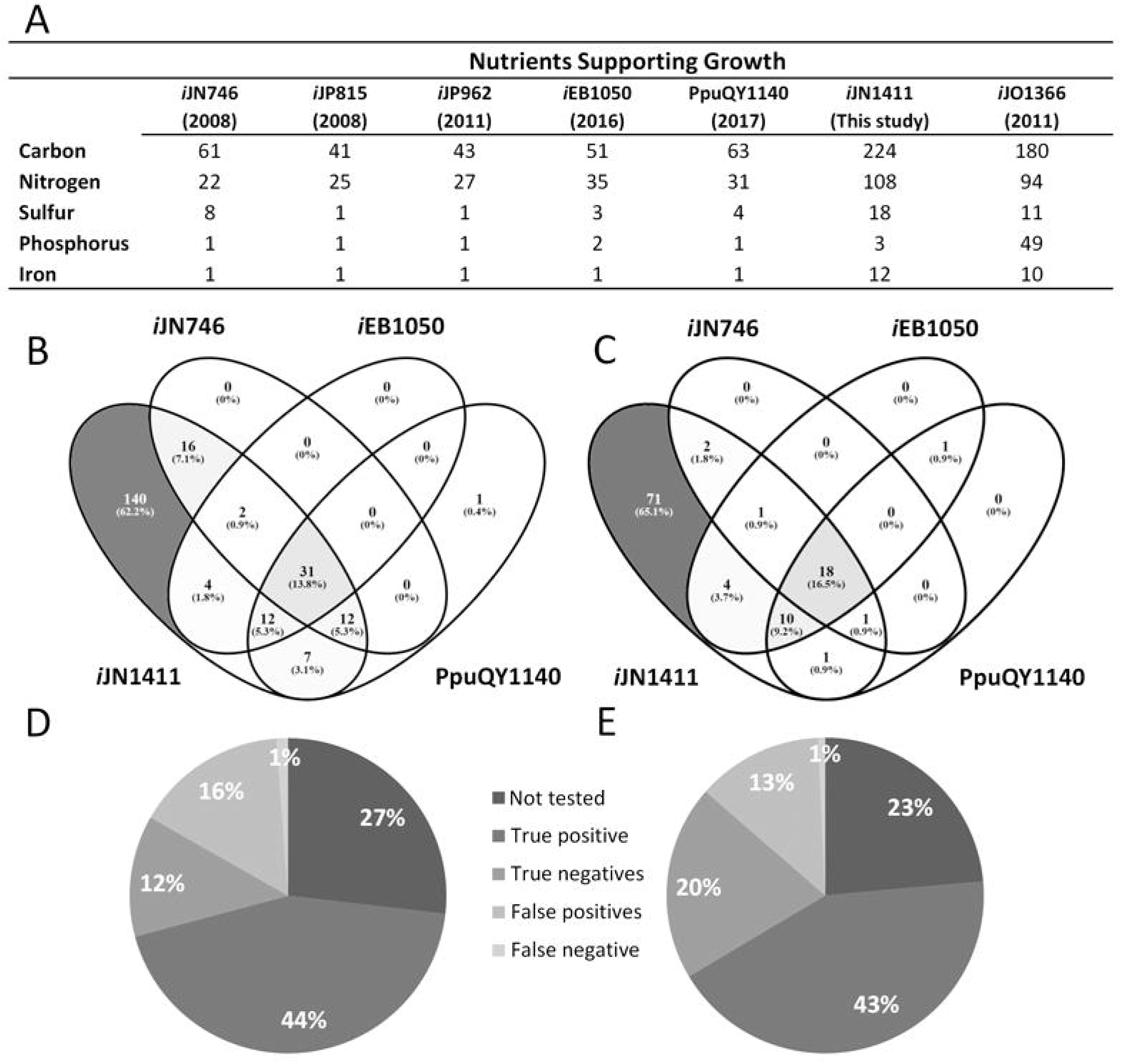
Identification and validation of nutrients supporting *i*JN1411 growth. **A.** The number of nutrients supporting growth in *i*JN1411, previous GEMs of *P. putida* and the latest GEM of *E. coli i*JO1366. **B and C.** A qualitative comparison of the carbon and nitrogen sources supporting growth in *i*JN1411, *i*JN746, *i*EB1050 and PpuQY1140. **D and E.** The prediction accuracy of *i*JN1411 for different carbon and nitrogen sources. Details are given in Table S2.

**Table 2.** Comparison of growth performance of *i*JN1411 with previous GEMs of *P. putida*. Constraints used are underlined. NA, not applicable. *i*EB1050 and PputQY1140 models lack of Octanoate and PHA metabolisms. For growth on octanoate as carbon source, nitrogen and oxygen uptake were constrained to 3.1 and 13.5 mmol.gDW^−1^.h^−1^, respectively.

| Carbon source | Uptake rate | Secretion rate (Gluconate)<br>(mmol.gDW<sup>-1</sup>.h<sup>-1</sup>) | Secretion rate (2-Ketogluconate)<br>(mmol.gDW<sup>-1</sup>.h<sup>-1</sup>) | Growth rate/PHA production rate (PHAC6 + PHAC8)<br>(h<sup>-1</sup>)/(mmol.gDW<sup>-1</sup>.h<sup>-1</sup>) |  |  |  |  | Reference |
| --- | --- | --- | --- | --- | --- | --- | --- | --- | --- |
|  |  |  |  | *iJN746* | *iEB1050* | *PpuQY1140* | *iJN1411* | *In vivo* |  |
| Gluconate | <u>5.1</u> | NA | NA | 0.58/NA | 0.67/NA | 0.37/NA | 0.47/NA | 0.43/NA | (65) |
| Glucose | <u>6.3</u> | NA | NA | 0.76/NA | 0.91/NA | 0.50/NA | 0.61/NA | 0.56/NA | (65) |
| Glucose | <u>7.3</u> | NA | NA | 0.86/NA | 1.05/NA | 0.59/NA | 0.71/NA | 0.73/NA | (52) |
| Glucose | <u>10.9</u> | <u>2.8</u> | <u>2.6</u> | 0.70/NA | 0.81/NA | 0.49/NA | 0.57/NA | 0.57/NA | (53) |
| Octanoate<sup>a</sup> | <u>3.4</u> | NA | NA | 0.31/1.9 | NA/NA | NA/NA | 0.2912/1.4<br>9 | 0.29/1.5 | (81) |

Because accurate predictions of growth rates alone cannot guarantee the quality of GEMs, we compared flux predictions on glucose to experimentally reported values (53). We found a good correlation between predicted and experimental values, with Kendall’s τ = 0.82, significantly higher than for the *i*EB1050 and PpuQY1140 models, τ = 0.53 and τ = 0.68, respectively (Fig. 3, Fig. S10). A well-known trait of *Pseudomonas* is the activation of the pyruvate shunt as a main source of oxaloacetate bypassing to the malate dehydrogenase (53, 54). Despite this alternative pathway being less efficient from an energetic point of view, this feature of *Pseudomonas* guarantees a high level of NADPH which is critical in order to provide metabolic robustness, including tolerance to oxidative stress (5, 53, 55). *i*JN1411, however fails to predict the activation of the pyruvate shunt as an alternative source of oxaloacetate that was somewhat expected since flux balance analysis (FBA) excludes suboptimal flux distributions (56). We therefore perform a sensitivity analysis of flux predictions as a function of the flux through Pyruvate Carboxylase (PC) (Fig. 2B). In good agreement with experimental results, the increasing of PC flux leads a large flux decreasing through Malate Dehydrogenase (MDH), a significant increase in the flux through Malic Enzyme (ME2) and a slight increase of TCA cycle, Pyruvate Dehydrogenase (PDH) and Pyruvate Kinase (PYK). When the experimental flux through pyruvate carboxylase was used as additional constraint, the accuracy in the flux distribution prediction increased significantly (τ = 0.98) (Fig. 3C). In summary, the flux predictions demonstrate the high accuracy of *i*JN1411, as well as the likely role of the mechanisms fueling metabolic robustness such as the Pyruvate Shunt, as one of the main mechanism disturbing the linearity of genotype-phenotype relationship (see below). *i*JN1411 can thus predict growth capabilities, growth rates and flux distributions for KT2440 with high accuracy, at a comparative level as the well-developed *E. coli* model does.

**Figure 3.**
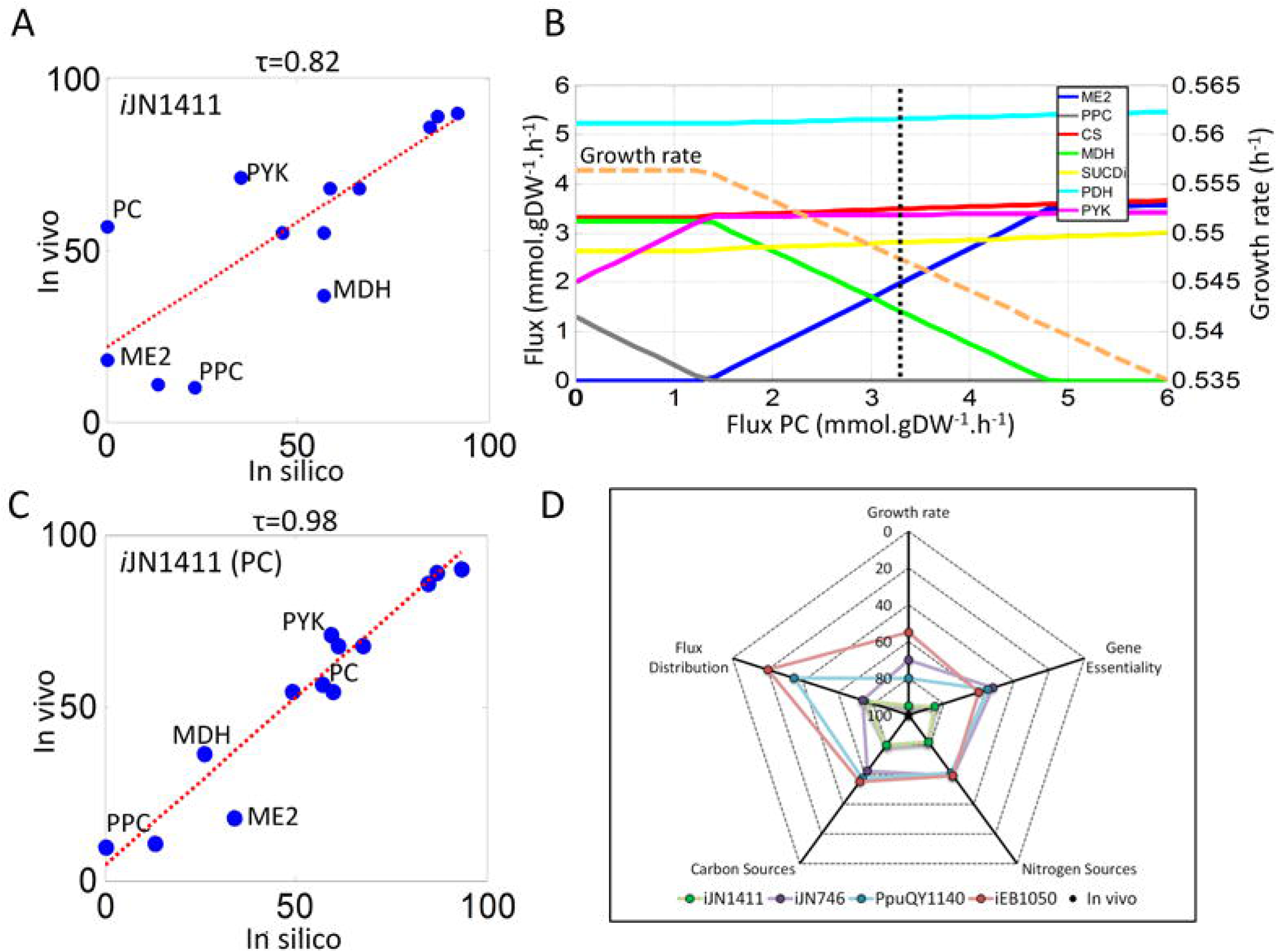
Validation of flux predictions and overall prediction accuracy. **A.** Comparisons between experimentally reported flux values in the central metabolism of *P. putida* growing on glucose (53) and predicted flux values obtained with *i*JN1411. **B.** Robustness analysis of flux predictions in *i*JN1411 obtained by varying the flux through Pyruvate Carboxykinase (PC), dotted line denotes reported flux for PC (3.24 mmol/gDW.h). **C.** The predicted values when the experimentally reported flux through the PC reaction was imposed as an additional constraint. Fluxes across the PC, Malic enzyme (ME2), Pyruvate dehydrogenase (PDH), Pyruvate kinase (PYK), Phosphoenolpyruvate carboxylase (PPC), Citrate synthase (CS) and Succinate dehydrogenase (SUCDi) are indicated. Correlation between *in vivo* and *in silico* flux values is expressed as Kendall’s rank correlation coefficient (τ). **D.** Perceptual accuracies of *P. putida* models with respect to in vivo data for growth rates, flux distribution, gene essentiality, carbon and nitrogen sources predictions are shown.

### Gene essentially data contextualization within *i*JN1411

The validation of GENREs through prediction of gene essentiality is a powerful way to assess and improve the accuracy of prediction while providing a suitable platform for the contextualization of knock-out mutant studies at the genome-scale (57–59). We performed a gene essentiality analysis on rich medium and then mapped the predicted essential genes with the knockouts available at Pseudomonas Reference Culture Collection (PRCC) (60). This approach defined an accurate *in silico* LB (*i*LB) medium and a core biomass objective function (See SI1, Table S3). A total of 117 essential genes were predicted under these conditions (Fig. 4, Table S4). The model was highly accurate in predicting essential genes. Only nine gene knockouts predicted as essential were found to be not essential in PRCC. These false positive essential genes were involved in the transport of cations and the biosynthesis of cofactors, suggesting alternative transport or biosynthetic mechanisms encoded in the genome of KT2440. The accuracy of *i*JN1411 was further evaluated on glucose minimal media against an experimental dataset (61). Up to 81 conditionally essential genes in glucose were predicted after excluding those also essential in *i*LB. Of those, 54 genes were present in PRCC and could be validated (Fig. 4C, Table S4). We found that *i*JN1411 was significantly more accurate than *i*JN746, *i*EB1050 and PpuQY1140 with 89% accuracy compared to 57%, 65% and 63% (two-sided p-values of Fisher‘s exact test was less than 10^−3^) respectively. The strain-specific BOF of *i*JN1411 allowed the correct prediction of several genes involved in cofactors biosynthesis as essential in contrast to previous reconstructions. In addition, *i*JN1411 was the only model able to predict the essentiality of the *edd* and *eda* genes, which encode key reactions of Entner-Doudoroff pathway, despite this pathway being well known to be essential for growth in glucose (54). Finally, the gene essentiality analysis provided a unique opportunity to gain new insights into the metabolism of KT2440. For instance, thought growth and gene knockouts analysis we prove the participation of the *cob* genes in the biosynthesis of vitamin B_12_, and we also show that they are only essential for the catabolism of a few nutrients such as ethanolamine (SI1, Fig. S11).

**Figure 4.**
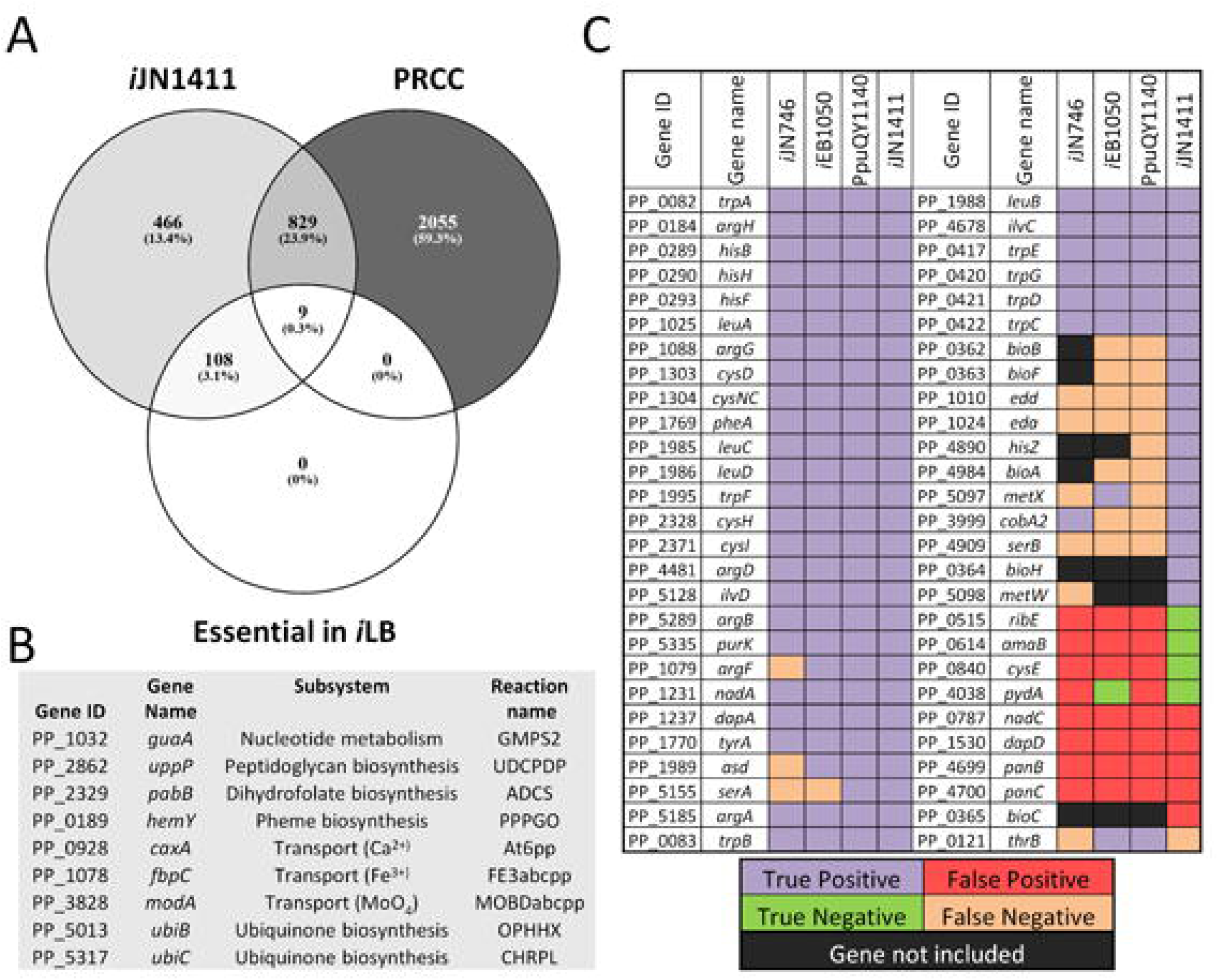
Gene essentiality analysis and validation. **A and B. Prediction of essential genes in *in silico* LB medium (*i*LB) and comparisons with experimental results.** Genes predicted to be essential in the *i*LB medium were compared with the gene content of *i*JN1411 and single gene knockouts present in the Pseudomonas Reference Culture Collection (PRCC) screened in rich medium. Only 9 false positives were predicted by *i*JN1411 and are shown in panel **B. C. The capabilities of *i*JN1411, *i*JN746, *i*EB1050 and PpuQY1140 for predicting essential genes in glucose minimal medium.** Purple and green denote genes that were correctly predicted as essential and non-essential, respectively, while red and orange denote the incorrectly predicted genes. Genes not included in the GENREs are shown in black.

### Functional assignment of metabolic capabilities of *P. putida* based on multi-strain modeling

Semi-automatic platforms for metabolic reconstruction are hampered by the lack of high-quality GENREs that can be used as templates for reconstructing phylogenetically related organisms. With the sole exception of the semiautomatic reconstruction of *Enterobacterias* which took advantage of a highly curated *E. coli* models (17, 19), this approach is still under-exploited. To show the potential of *i*JN1411 as template for modeling *Pseudomonas* group we performed a reconstruction of the all *P. putida* strains sequenced to date by employing similar approach to the one previously used for modeling *E. coli* and *Shigella* strains (17) (See Methods, Table S5). This approach resulted in highly complete metabolic models, which share more than 95% of the reactions included in *i*JN1411 (Fig. 5). Furthermore, by keeping only those genes present in all the *P. putida* strains, a core-genome metabolic model of *P. putida* (PP_CORE) was obtained. This model possesses only the common metabolic capabilities of all the sequenced strains of this species.

**Figure 5.**
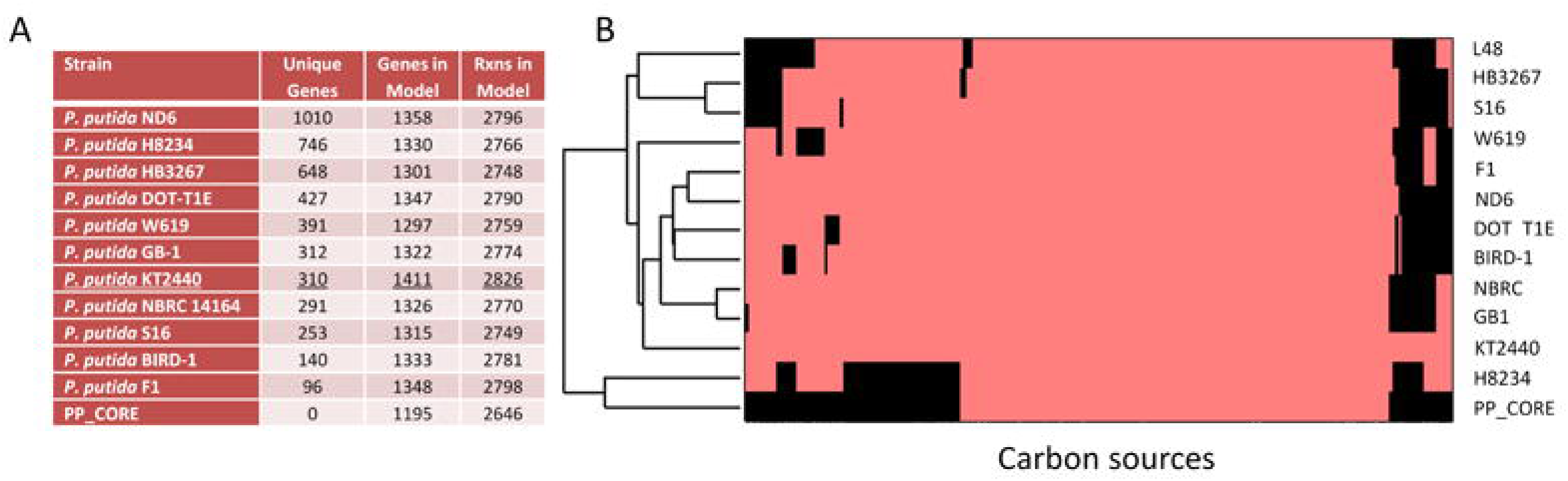
Genome-scale modeling of the *P. putida* group. **A. Metabolic content of *P. putida* GEMs.** The table summarizes the number of genes and reactions in each GENRE. The number of unique genes in each strain is indicated. **B. Metabolic versatility of *P. putida*.** The metabolic versatility of each *P. putida* strain was estimated by maximizing growth of the corresponding GEM using the 223 carbon sources supporting growth in *i*JN1411. In pink are those carbon sources supporting growth in each GEM while black indicates the absence of growth.

Finally, we evaluated the metabolic capabilities included in each model by analyzing the array of carbon sources supporting growth (Fig. 5B). We found that the strain-specific models largely shared the high metabolic versatility of *i*JN1411. H8234 stood out as the strain with the lowest metabolic versatility. Interestingly, this strain was isolated from a hospital patient presenting with bacteremia, which could explain the loss of metabolic capabilities when compared with environmental isolates (62). Overall, our analysis shows that metabolic versatility is a general feature of the *P. putida* group, irrespective of the strain.

### De-composition of metabolic robustness in *P. putida*

Despite metabolic robustness being one of the main features of *P. putida* (34, 52, 63), the molecular mechanisms fueling this emergent property are poorly understood, and only recently genomics approaches have brought some light on this issue (30). In order to address the breakdown of the metabolic robustness of KT2440, at the genome scale, we performed a gene essentiality analysis using *i*JN1411 in a set of 385 different environmental conditions including alternative sources for carbon, nitrogen, sulfur, phosphorous, and iron, as well as the exposure to heavy metals and stressors.

This analysis revealed that KT2440 has a high buffering capability against genetic and environmental perturbation, as only 106 genes (7.5% of the genes in *i*JN1411) were found to be essential in all the conditions analyzed (SEG) (Fig. 6, Table S6). The average number of essential genes per condition was around 200 (≈14%), with LB rich medium being the condition with lower number of essential genes, 124. A total of 501 genes were essential in at least one condition. This correspond to a surprisingly high number of genes being non-essential, 910 (65%), in any condition.

**Figure 6.**
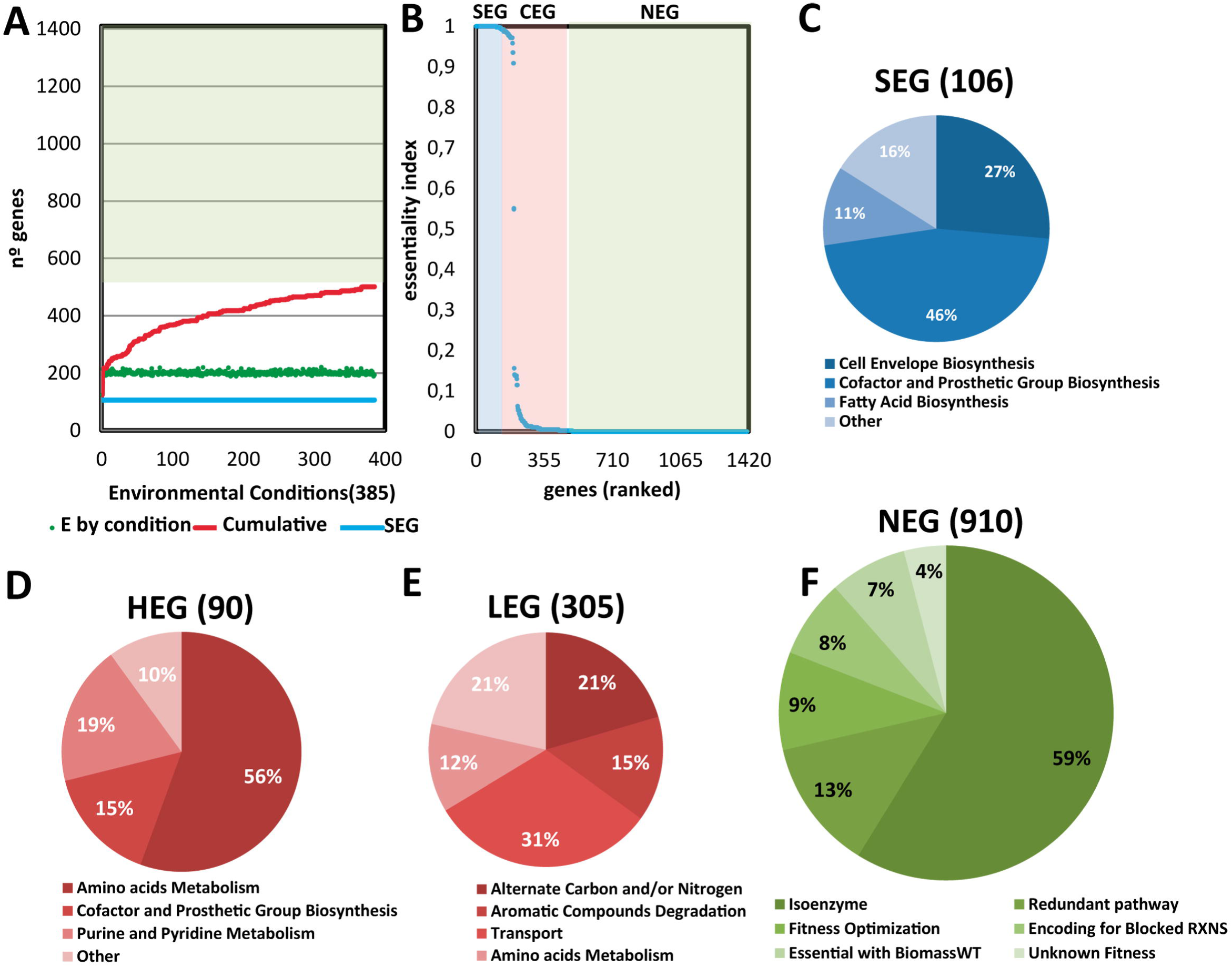
Decomposition of the metabolic robustness of *P. putida* under environmental and genetic perturbations. **A.** The genes from *i*JN1411 were deleted sequentially and the growth rate of the resulting *in silico* knockout strains computed in 385 different environmental conditions. The number of essential genes (EG) in each condition, the cumulative number of essential genes and the number of genes essential in all conditions (super-essential genes, SEG) are shown. **B.** A graphical representation of the essentiality index (ei) of each gene in the reconstruction. The ei is defined as the number of conditions in which a target gene was predicted to be essential divided by the total number of conditions simulated. **C, D** and **E.** <u>T</u>he distribution over subsystems of predicted super-essential genes, highly essential (HEG, 0.5<ei<1) and low essential genes (LEG, 0<ei≤0.5) respectively. **F.** A breakdown of genes predicted as non-essential across all conditions (NEG) in terms of function.

We further calculate a measure of gene essentiality for each gene in the reconstruction. The essentiality index (*ei*) is defined as the number of conditions in which a gene was predicted to be essential divided by the total number of conditions simulated. We then grouped the genes included in the reconstruction into three categories, i) genes that were essential in all the conditions tested (SEG, *ei*=1), ii) genes non-essential in any condition (NEG, *ei*=0), and iii) conditional essential genes (CEG) which were essential in at least one condition (0<*ei*<1). The 395 CEG were additionally grouped as high essential genes (HEG 0.5<ei<1) and low essential genes (LEG 0<ei<0.5), see Fig. 6.

The analysis of the SEG genes showed that they were restricted to the *P. putida* core genome (Table S6) (30), and mainly confined to anabolism e.g., biosynthesis of cofactor and prosthetic groups and cell envelope (Fig. 6). No catabolic processes were identified as super-essential in KT2440. These results are in good agreement with recent predictions of essential reactions in biological networks which showed a high grade of super-essentiality of anabolic reactions in nature over key catabolic processes such as central carbon metabolism (64).

The genes classified as HEG and LEG differ considerably in the metabolic processes in which they are involved. Thus, while the 75% of the genes classified as HEG were responsible for the biosynthesis of amino acids and nucleotide metabolism, the LEG genes mainly provide metabolic versatility to KT2440 including subsystems such as alternate carbon and/or nitrogen sources, degradation of aromatic compounds, transport and amino acids catabolism. The NEG were widely distributed along the subsystems present in *i*JN1411 (Table S6).

When we investigated the reason behind of the lack of essentiality of NEG, we found that around 75% of them provide robustness under genetics perturbations through the alternative (redundant) mechanism. Thereby, 59% of the NEG have at least one isozyme which could replace each other. Similarly, we also found redundant metabolic pathways for 13% of the genes classified as NEG. This trait is well-known in KT2440; for instance, up to 3 different peripheral pathways have been reported for the initial catabolism of glucose (65) and lysine (66). Another important group of NEG was involved in growth optimization since their absences either decreased the growth rate (fitness optimization) or they were only essential with respect to the complete biomass function. This large number of non-essential genes in *P. putida* is in line with gene essentiality analysis in robust generalist bacteria such as *E. coli*, 93% (67) or *P. aeruginosa*, 94% (67, 68), but contrast with that reported in specialist organisms such as cyanobacteria (69).

### Systematic identification of ATP-fuelled metabolic robustness modules in *P. putida*

The fast-growing use of metabolic flux analysis to study bacterial physiology (70), when combined with the *in silico* flux analysis provided by genome-scale models (71), have revealed the nonlinear relationship between phenotype and genotype in terms of flux, and often, the *in silico* predictions differ from those experimentally determined (72). It has recently been shown via the application of multi-objective optimization that flux distributions computed using metabolic models agree with experimentally determined values when a combination of maximum ATP yield, maximum biomass yield, and flux adjustment between multiple environmental conditions were used (72). This excess of ATP production could increase biological robustness by acting as biological fuel towards unexpected perturbations at the expense of lower fitness under stable conditions. Therefore, it is reasonable to assume that metabolic robustness is a key systems emergent property disturbing the linearity in the genotype-phenotype relationship.

The molecular mechanisms enabling overproduction of ATP are less clear since its storage by living systems, under the assumption of a biological steady state, is limited. In order to maintain an elevated production of ATP, a likely strategy is the development of mechanisms allowing proper turnover of ATP. Energy-dissipating futile cycles based on enzymes involving phosphorylation and dephosphorylation or transport systems are well-known in living systems (73, 74). They are dependent on the physiological state and act to balance the ATP/ADP ratio when growth is limited by nutrients other than energy, however their participation is unlikely under optimal growth conditions. Therefore, we became interested in investigating whether some of non-essential metabolic genes in *i*JN1411 could be involved in balancing the ATP/ADP ratio in a biological steady state. This was done by identifying ATP-dependent cycles in the network by applying an enumeration algorithm (75). We identified 337 reactions taking part in 544 futile cycles composed of at least two reactions (Table S7). We excluded from the analysis futile cycles based on transport reactions and coupled kinases/phosphatases since we were not interested in studying conventional futile cycles (74, 75). A large number of cycles included a common core of reactions, thus it was possible to reduce even more the number of potential futile cycles based on this common core of reactions (Table S7). Finally, we applied two additional criteria in order to identify putative metabolic cycles involved in ATP-fueled metabolic robustness: i) a non-zero flux through the cycle should lead to a reduction in the growth rate, and ii) the reactions forming part of the cycle should be encoded by NEG genes. Nine cycles fulfilling these criteria were identified: cycles of pyruvate, oxalacetate, glutamate, polyphosphate (Poly-P), trehalose, glycogen, fatty acids, PHA, and PRPP (Fig. 7).

**Figure 7.**
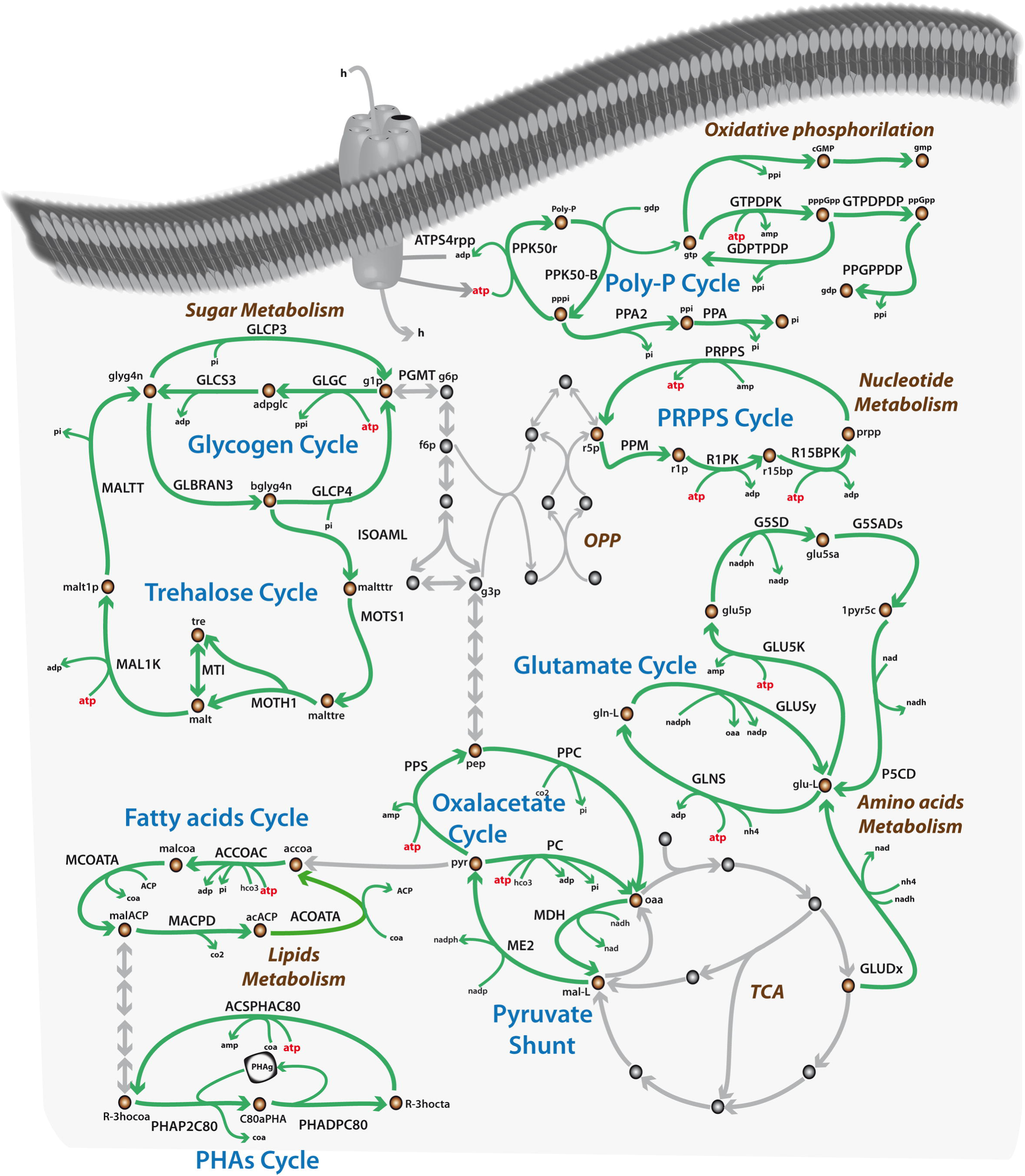
Graphical representation of ATP-fueled metabolic robustness cycles in *P. putida.* The ATP-consuming cycles providing metabolic robustness to *P. putida* are shown in green and blue while the main elements of the primary metabolism fed by such cycles, i.e. the metabolism of lipids, amino acids, sugars, nucleotide, the TCA cycle and oxidative phosphorylation are indicated in brown. The ATP consumed is shown in red. The abbreviations for reactions and metabolites are given in Table S1.

In addition to the expected energy-dissipating properties of these cycles, we noted that they additionally provide an ATP-fueled mass-balanced flow of metabolites around key metabolic nodes (Fig. 7). Thus, the glutamate cycle provides flow of nitrogen metabolites around the amino acid metabolism and the pyruvate and oxaloacetate cycles (pyruvate shunt) recirculate organic acids around the TCA. The fatty acids and PHA cycles keep moving fatty acids-like compounds around and the glycogen and trehalose cycles provide an effective turnover of sugars and phosphosugars. Finally, the Poly-P cycle supports the turnover of deoxynucleoside triphosphate around oxidative phosphorylation and the PRPP cycle drives the flow of pentose phosphate around the nucleotide metabolism. Of the 36 genes encoding these cycles, 34 belong to the core genome (30), supporting the idea that the presence of these ATP-fueled metabolic cycles is a conserved feature in *P. putida*, as species.

Following this hypothesis, and in contrast to conventional futile cycles which are solely expressed under stress conditions, the genes encoding these newly identified metabolic cycles should be highly expressed under exponential growth phase, irrespective of the nutritional scenario. By using gene expression datasets from *P. putida* KT2440 obtained in exponential growth phase (76), we proceed to analyze if this was indeed the case. To complete the analysis we performed a gene essentiality analysis using *i*JN1411 under the four carbon sources for which transcriptomic data were available, e.g., glucose, fructose, glycerol, and succinate (Table S8, Fig. S13). This enables the comparison of gene expression levels between essential and non-essential genes in each nutritional condition. As expected, we found higher expression levels and lower data variability in the genes predicted to be essential in all the conditions. For instance, while the average expression level of essential genes in glucose was 450 reads per kilo base per million mapped reads (RPKM), non-essential genes had a significantly lower value, 189 RPKM.

We further focused on the expression level of genes encoding enzymes participating in ATP-fueled metabolic cycles in order to determine if they were expressed. We found that they exhibited gene expression values even higher than essential genes (Fig. 8, Table S8). However, since bacterial gene expression is continuous, the stablishing of a threshold gene expression value to consider a gene to be significantly expressed is uncertain. A well-accepted assumption is that which consider a given gene expressed if its expression level is under the 25th percentile of the expression data (77). To support further this assumption, we searched for the expression level of genes experimentally seen as not significantly expressed in these conditions (Table S8), among others the *gal genes* (PP_2513-9) involved in the degradation of gallate (78). The expression level of these genes was around 10 RPKM and always within the 25th percentile in all the conditions (Table S8) that supports the notion that the genes encoding ATP-fueled metabolic cycles are, indeed, not only expressed but highly expressed despite the negative impact on growth rate.

**Figure 8.**
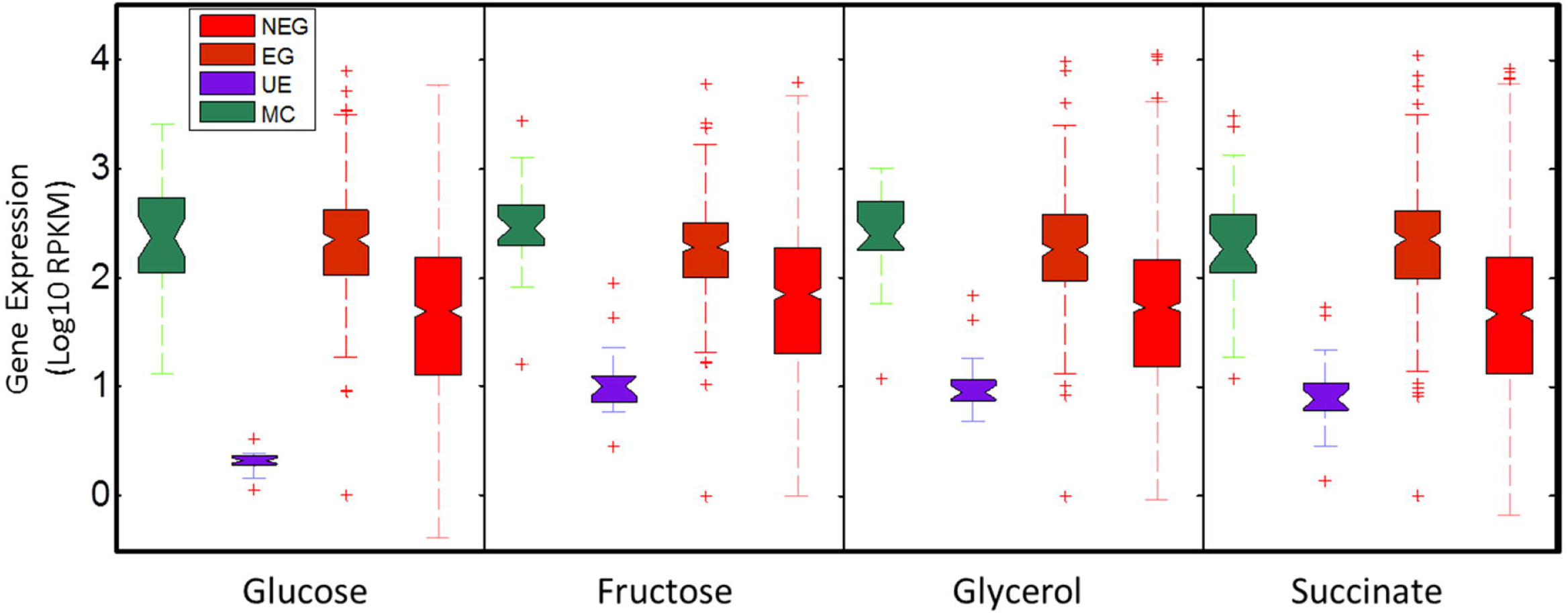
Contextualization of the genes encoding for robustness cycles in *P. putida* with respect to gene essentiality and gene expression. Box plots of gene expression values of each of the groups identified above. The edges of the boxes represent the 25^th^ and 75^th^ percentage, respectively while the midpoints represent the median expression values. The notches represent comparison intervals. Two medians differ significantly at the 5% level if their intervals (notches) do not overlap. The upper and lower parts of the whiskers represent the maximum and minimum gene expression values. Outliers are indicated by red crosses.

Taken together, the cyclic nature, and the level of expression of these ATP-fueled metabolic cycles under exponential growth phase, which exclude any potential metabolic unbalance, leads us to think that such cycles are not conventional futile cycles but buffering cycles. In other words, these cycles would be acting keeping key metabolites recirculating around central metabolism providing stability in fitness, thus providing metabolic robustness under changing environmental conditions. The operability of these cycles in robust generalist bacteria, such as *P. putida*, would inevitably contribute to disturb the linearity of genotype-phenotype relationship in term of carbon flux distribution. Supporting this idea, we have shown above how the flux through the pyruvate shunt is the main contributing factor for the *in vivo*/*in silico* flux distribution discrepancy when glucose is used as the sole carbon and energy source (Fig. 3).

## Discussion

### *i*JN1411 expands the metabolic reactome available for computation

A detailed metabolic model is a powerful tool for analyzing the systems metabolic properties of its target organism (21, 22, 69). The level of completeness and accuracy of *i*JN1411 makes it one of the largest and high-quality genome-scale reconstructions built to date. The careful reconstruction process allowed detailed modeling of *P. putida* catabolism and anabolism beyond that of what was known (Fig. 2). *i*JN1411 expands the metabolic reactome available for computation, including the metabolic signatures of *Pseudomonas*, a bacterial group with significant biotechnological and clinical interest (24, 34). The comparison of *i*JN1411 predictions with experimental data shows a high level of accuracy compared to previous models (Fig. 3D). Despite the large effort done on the metabolic reconstruction of *P. putida*, the current GEMs, exhibit similar performance to *i*JN746 (Fig. 3D), the first GEMs published almost a decade ago, in terms of i) computable reactome (active reactions) (Fig.1, Table 1), ii) nutrients supporting growth (Fig.2), iii) growth rate predictions (Table 2), iv) carbon flux predictions (Fig.3) and v) gene essentiality predictions (Fig. 4). Mapping the available computable reactome for a single strain is useful to detect inconsistences between different GEMs, but this approach, itself, is not enough for constructing high-quality GEMs. Instead, it should be used as a preliminary step followed of a careful manual curation and experimental validation process, as we have done here. As we previously warned (13), an abusive use of the so-called consensus approach for modeling without the proper validation and experimental contextualization could lead the inclusion of inaccurate metabolic content, closer to be a collage rather than strain-specific.

In contrast, *i*JN1411 has proven to be a useful tool for reconstructing other *P. putida* strains. The functional comparison between these strains highlighted that the metabolic versatility and robustness are metabolic traits inherent to the whole *P. putida* group. Nevertheless, this collection of *P. putida* strains GEMs should be considered as drafts, which still require of careful manual curation, work that is currently ongoing in our lab. Finally, because an increasing number of *P. putida* strains have been and continue to be isolated for strain-specific biotechnological and/or bioremediation purposes, the reconstruction of a *P. putida* core genome-scale model has great biotechnological potential. The PP_CORE model can be widely used for the computational analysis of a huge amount of biotechnological applications curried up by *P. putida* strains whose genomes have not been sequenced yet, simply by adding the reactions responsible for such processes.

### Bacterial metabolic robustness capacitators

A significant number of genes non-essential for growth were found to participate in cycles consuming ATP, irrespective of their primary metabolic function (Fig. 6–7). Futile cycles occur in micro-organisms inducing a considerable energy burden for the cell (74, 75, 79, 80). These cycles operate when reactions act in an antagonistic fashion, simultaneously promoting the dissipation of energy. Overall, such cycles fall in two main categories, those involving simultaneous phosphorylation and dephosphorylation reactions and those involving transport reactions in the opposite direction (74). Therefore, it has been suggested that energy spilling reactions are i) a common feature of growth with an excess of energy, and ii) an indicator of the imbalance between anabolism and catabolism (73). For instance, we recently showed how cyanobacterial photosynthetic networks activate a large array of energy dissipating mechanisms for balancing the ATP/NADPH ratio under carbon limitation and/or high light conditions (69).

We identified in our analysis hundreds of putative conventional futile cycles, e.g., coupled kinase and phosphatase and coupled transport reactions (Table S7). However, we additionally identified a set of atypical futile cycles providing a mass balanced flow of metabolites around basic metabolic hubs. In contrast to conventional futile cycles, we found that the genes encoding these cycles were highly expressed in the exponential growth phase, suggesting they are not induced by nutrient limitation and/or imbalanced metabolism. To the best of our knowledge, there are not systematic experimental studies focused on the functionality and biological role of these ATP-fuelled metabolic cycles. However, several lines of evidence support many of them being active in *Pseudomonas* and playing an important biological role. For instance, it is known that a large amount of the TCA cycle carbon flux occurs through the so-called pyruvate shunt, which bypasses the conventional and more energetic pathway through malate dehydrogenase (Fig. 3) (65). This alternatively provides a higher level of NADPH that is required for fueling mechanisms against oxidative stress (5, 53, 55). Here we show that flux through pyruvate shunt is primary responsible of disturbing the linearity of phenotype-genotype relationship in *P. putida* on glucose. This behavior decreased growth rate slightly (0,556 vs 0,547 h^−1^), but provided additional NADPH equivalents to face unexpected and sudden environmental insults. It should therefore be expected that other robustness cycles can replace pyruvate shunt under different environmental conditions. For instance, the cyclic nature of PHA metabolism and its function as a futile cycle dissipating energy has been suggested in *P. putida* when fatty acids are used as carbon sources (81, 82). The fact that the PHA cycle provides more robust growth during transient nutrient conditions lends further supports our hypothesis suggesting that such cycles, indeed, can act as sources of biosynthetic building blocks and energy under environmental perturbations. Finally, it is noteworthy that *P. putida* strains lacking the Poly-P cycle exhibit a large lag growth phase (83), suggesting a key role of this cycle as a guarantor of fitness under changing environmental conditions. We are not aware of any experimental validation of the rest of metabolic cycles identified in our analysis, but based on gene expression data and their cyclic nature they could be expected to have a similar function.

In summary, our computational analysis supports the idea that, far from being futile cycles, the ATP-fueled metabolic cycles identified could act as buffering cycles or “metabolic capacitors” surrounding the primary metabolism and providing a pre-processed source of energy and anabolic building blocks while balancing and optimizing the redox state. Additionally, the operation of such cycles *in vivo* at expense of ATP agree with the suggested multi-objective of bacterial networks (72) and could explain, to a great extent, the nonlinear genotype-phenotype relationship, in terms of flux distributions, as pyruvate shunt does on glucose catabolism.

#### Updating the metabolism’s structure of robust environmental bacteria

Similar to other bacteria, the metabolism of *P. putida* follows the so-called “bow tie” model, where nutrients are catabolized along a catabolic funnel to produce the precursors and energy required for synthesis of building blocks (84, 85). Thus, the bow tie can be decomposed into three modules, catabolism and anabolism, which are organized as the fan-in and fan-out part of the bow tie, and the knot, which includes the central metabolism. Bow tie structure of metabolism thus facilitates robust biological functionality. Recently, Sudarsan and colleagues provide strong evidence suggesting two different operational modes in the *P. putida* bow tie. While catabolism and anabolism are shown to be highly flexible and robust with a large correlation between metabolic flux and transcriptional levels, the central metabolism was extremely stable, showing no correlation between metabolic fluxes and transcriptional expression. As a result, it was suggested that the central metabolism of *P. putida* is finely regulated at the posttranscriptional and metabolic levels (85). In light of our metabolic analysis and based on the proposed bow tie model, two interesting questions arise: i) how do bacteria increase metabolic robustness under this metabolic structure, and ii) how do bacteria merge the highly flexible catabolism and anabolism modules to the rigid and stable central metabolism?

Regarding the first question, the large metabolic versatility found in *P. putida* is noteworthy. Among the carbon and nitrogen sources supporting growth we found key anabolic precursors, including amino acids, sugars, fatty acids, nucleotides. This trait provides high robustness against genetic and nutritional perturbation by avoiding potential deleterious effects due to mutations in biosynthetic pathways and/or nutrients depletion. *P. putida* also exhibits a versatile anabolism, providing a large array of mechanisms to modify macromolecules and/or synthesize *de novo* new ones in response to environmental changes. For instance, under water-limiting conditions, *P. putida* produces alginates in order to maintain a hydrated microenvironment, thus protecting itself from desiccation stress and increasing its chances of survival (86). Thus, it is reasonable to think that robust bacteria such as *P. putida* increase metabolic robustness by expanding the arsenal of both, catabolic and anabolic pathways (Fig. 9). In addition, the presence of redundant isozymes and/or metabolic pathways was shown to be responsible for increasing genetic robustness in *P. putida*. Therefore, the bow tie model should be understood as multidimensional in robust bacteria including several functional redundant metabolic layers, thus contributing even more to increased metabolic robustness (Fig. 9).

**Figure 9.**
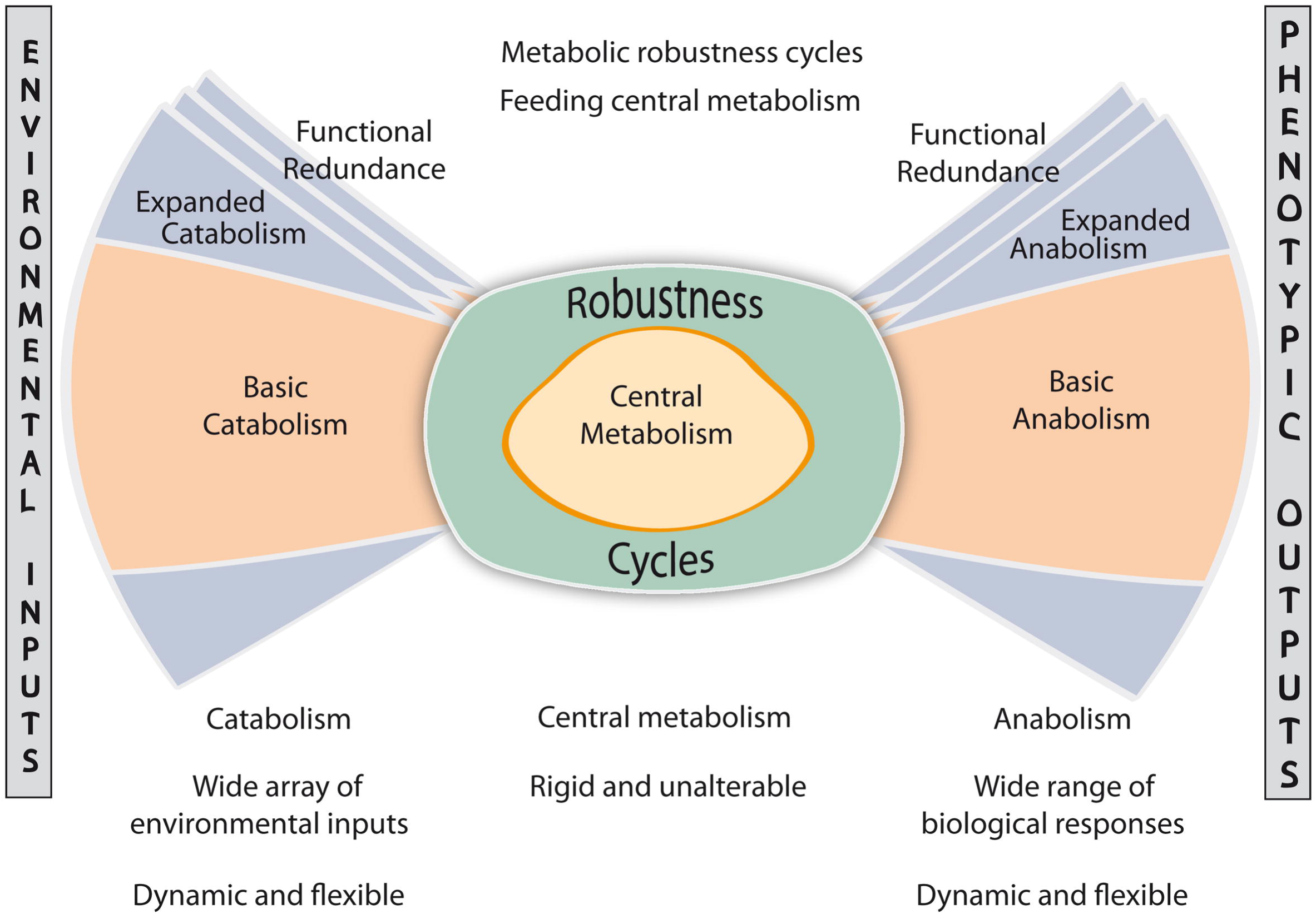
The bow-tie structure of the *P. putida* metabolism. A metabolic network can increase its metabolic robustness by: i) expanding catabolism and anabolism; ii) increasing functional redundancy for many metabolic processes; and iii) incorporating buffering cycles around central metabolism (metabolic robustness cycles), acting as metabolic capacitors to promote the stable connection of catabolism and anabolism with the central metabolism under perturbations.

As to the second question, it is tempting to think that the bow tie model is still incomplete and that additional metabolic mechanisms allow the stable transition from catabolism and anabolism to central metabolism, irrespective of the environmental inputs. The ATP-fueled metabolic cycles identified in our analysis could indeed fulfilling this task be acting as “metabolic capacitors” (Fig. 9). They would provide a constitutive flow of key metabolites around the central metabolism independent of the nutritional conditions. This carbon flow could feed transitory central metabolism under perturbations such as nutrient depletion and environmental insults protecting the optimal functionality of the central metabolism while avoiding the requirement of large changes on gene expression (Fig. 9).

In summary, we here present a high-quality metabolic modeling of *P. putida* which represents a large expansion of the current computable metabolic space including important modules beyond primary metabolism. The systematic and contextualized analysis of this new metabolic space revealed its role in disturbing the genotype-phenotype relationship. Furthermore, we have shown how this non-essential metabolism for growth plays a key role in the metabolic robustness in *P. putida*, paving the way for i) a better understanding of the genotype-phenotype relationship, ii) engineering of metabolic robustness cycles in biotechnology and iii) identifying potential drug targets in robust pathogens such as *P. aeruginosa*.

## Material and Methods

### Metabolic reconstruction process of *P. putida* KT2440

The workflow of the reconstruction process started with the GEMs of *P. putida* available at that time (November 2011): *i*JN746 (36), *i*JP850 (37), PpuMBEL1071 (38), and *i*JP962 (39). As often happens for bacteria being reconstructed, the available models for *P. putida* KT2440 are significantly different and surprisingly they only share 523 genes from the 1213 unique genes included in the four models. This data highlights the large bias and lack of manual curation inherent in many metabolic reconstruction processes (13, 14). The overall workflow for the reconstruction process used here is shown in the supplemental material (SI1, Fig. S1). Briefly, the reconstruction was performed manually following an iterative tri-dimensional expansion based on genome annotation, biochemical, and phenotypic legacy knowledge. For genome-based expansion, the *P. putida* KT2440 model, Seed160488.1, (PputSEED) was downloaded from the Model SEED database (87) and its content mapped on *i*JN746. The subsequent comparison shows that PputSEED included 497, 881 and 655 exclusive genes, reactions and metabolites, respectively (SI1, Fig. S2, Table S1). These sets of genes, reactions and metabolites absent in *i*JN746 were further manually investigated one by one in order to justify their inclusion in the updated model based on legacy knowledge and/or computational evidence. When appropriate, the new content was included in the reconstruction and the new reactions and metabolites were named following the BIGG nomenclature (88). This approach significantly increased the content of *i*JN746, however we found several inconsistences in PputSEED in four categories; i) lack of specie-specific reaction formulations, including inappropriate substrate and/or cofactors and/or reversibility; ii) inaccurate GPR associations; iii) inaccurate compartmentalization of reactions; and iv) incorrect modeling of most of the biosynthetic pathways including the cell envelope, phospholipids. Multiple reactions were therefore excluded or reformulated based on *Pseudomonas* legacy knowledge.

In the second step, the content from two additional metabolic reconstructions of *P. putida*, *i*JP962 (39) and PpuMBEL1071 (38) were investigated following the above workflow and when appropriate, new content was added to the model. This second genome-based expansion step provided minor additions compared to PputSEED and similarly, a large number of genes were discarded and/or included in different GPRs. Following the recommendations transparence guidelines for metabolic reconstructions (14), the list of discarded genes from previous GENREs of *P. putida* (up to 336) and the reason of their exclusion are provided in Table S1.

The biochemical and phenotypic expansion was performed simultaneously by modeling new anabolic and catabolic pathways (SI1, Fig. S1). During this step, legacy knowledge from *Pseudomonas* found in databases such as the Pseudomonas Database, BRENDA etc., as well as in the primary literature was widely used. As a result, up to 409 unique citations are included in the final reconstruction and 2035 of the reactions have, at least, one citation supporting its inclusion in the reconstruction. The list of citations is provided in SI (Table S1). This detailed search for biological knowledge in *Pseudomonas* beyond the genome-annotation allowed the accurate modeling of multiple new biosynthetic and catabolic pathways, many of which were previously unknown in *P. putida* KT2440 and have been modeled here for the first time. Finally, the model built on the reconstruction was thoroughly evaluated in order to detect inconsistencies by experimental nutrient phenotyping and gene essentially data (60, 61) (SI, Table S2). This approach allowed the re-annotation and/or more accurate assignment of function to 246 genes encoded in the *P. putida* KT2440 genome. The complete list of re-annotated genes is provided in Table S1.

The SimPheny™ (Genomatica Inc., San Diego, CA) software platform was used to build the reconstruction. All the metabolites in the reconstruction were introduced according to their chemical formula and charge using their pKa value at pH 7.2. All reactions were subsequently mass and charge balanced. The reversibility for each reaction in the reconstruction was determined from the primary literature, when possible, or taken for phylogenetically related organisms. In addition, for each reaction included in the model a confidence score (CS) ranging from 1 to 4 was assigned (Table S1). A value of 1 indicates *in silico* evidence supporting the inclusion of a given reaction, e.g., the reaction is solely required for the functionality of the model. A value of 2 indicates genomic or physiological evidence. Reactions with a confidence score of 3 are supported by genetic evidence such as knockout characterization and a value of 4 indicates that the target GPR has been completely characterized. The average CS was 2.59 (Table S1). The model in SBML format is provided in SI3 and it will be made available in the BIGG database after publication.

### Constraints-based analysis

The *i*JN1411 model was exported from SimPheny as an SBML file and analyzed with the COBRA Toolbox v2.0 (89) within the MATLAB environment (The MathWorks Inc.). Tomlab CPLEX and the GNU Linear Programming Kit (http://www.gnu.org/software/glpk) were used for solving the linear programing problems. The constrain-based model consists of a 2087 × 2826 matrix containing all the stoichiometric coefficients in the model of 2087 metabolites and 2826 reactions (**S**). Flux balance analysis (FBA) was used to predict growth and flux distributions (56). FBA is based on solving a linear optimization problem by maximizing or minimizing a given objective function **Z** subject to a set of constraints. The constraints **S·v = 0** correspond to a situation of steady-state mass conservation where the change in concentration of the metabolites as a function of time is zero. The vector **v** represents the individual flux values for each reaction. These fluxes are further constrained by defining lower and upper limits for flux values. For reversible reactions an upper and lower bound of −1000 mmol.gDW^−1^.h^−1^ and 1000 mmol.gDW^−1^.h^−1^ were used, respectively. A lower bound of 0 mmol.gDW^−1^.h^−1^ was used in case of irreversible reactions. For simulating condition-specific growth conditions, lower bounds of the corresponding exchange reactions were modified accordingly (See SI1). By default, the maximum growth rate was used as the cellular objective. Additional model constraints sink and demand reactions required for the functionality of the model can be found in SI1.

For modelling and analysis some additional constraints were applied. The bounds of the Pit7pp (Na-dependent phosphate transport) reaction were constrained to 0 mmol.gDW^−1^.h^−1^ to avoid unrealistic ATP production. Sink and demand reactions are modeling reactions required for the functionability of the model. Sink reactions are included in order to provide key metabolites of unknown origin while demand reactions are required for the removal dead end metabolites. *i*JN1411 includes two sink reactions, sink_PHAg and sink_pqqA which provide the PHA granule required for PHA biosynthesis and the initial peptide required for PQQ biosynthesis, respectively. There are 31 demand reactions of which six are required to allow dead metabolites to leave the system e.g., DM_acmum6p, DM_5DRIB, DM_acgam, DM_AMOB, DM_doxopa, and DM_tripeptide, and 25 are needed for allow the accumulation of cytoplasmic polymers, including the 24 monomers of PHA and polyphosphate

#### Expansion and validation of nutrient sources supporting growth

To model the metabolic versatility of *P. putida* KT2440 primary literature and high-throughput nutrient phenotyping analyses of *Pseudomonas* spp were extensively scrutinized. The identification of a nutrient supporting growth in any *Pseudomonas* spp was the starting point for searching for potential genes encoding this ability in the genome of KT2440. If enough computational evidences supported the inclusion of the target catabolic pathway based on sequence identity, the corresponding reactions were added to the reconstruction. This iterative process resulted in the inclusion of hundreds of new reactions, many of them modeled in *i*JN1411 for the first time. This process concluded by adding the corresponding transport and exchange reactions. The transport reactions databases TCDB (90) and TransportDT (91) were used for this purpose.

The potential nutrients sources supporting growth on *in silico* M9 medium (*i*M9, See SI1) including glucose, ammonium, inorganic phosphate, sulfate and Fe^2+^ as default carbon, nitrogen, phosphate, sulfur and iron sources, respectively were identified *in silico* by maximizing the BOF. Carbon sources were identified constraining the glucose uptake rate to zero and testing sequentially all the metabolites for growth which an exchange reaction was present in the reconstruction. Nitrogen, sulfur, phosphate and iron sources were predicted similarly by constraining the uptake of the corresponding default nutrient to zero. Any metabolite providing a non-zero growth rate was considered as true nutrient.

The predicted carbon and nitrogen carbon sources were subject to bibliomic and/or experimental validation. All disagreements between predicted and experimental values were further carefully analyzed. Several false negatives (growth in vivo, but not in silico) were resolved by manual gap-filling resulting in the inclusion of new reactions and genes in the reconstruction. This process contributed to the reannotation of many metabolic genes in *P. putida* (the annotation update can be found in SI1, Table S1). If the gene encoding the target enzymatic activity was unknown, we decided to fit the experimental data by including orphan reactions only if enough bibliomic support was available. For instance, while the gene encoding the coniferyl alcohol dehydrogenase (COALCDH) appears to be missing from the genome of the KT2440, coniferyl alcohol can be utilized as the sole carbon and energy source by this strain (42). For false positives (growth in silico, but not in vivo), the criteria that we followed was to keep the corresponding catabolic pathway in the model if strong computational evidence (sequence identity) was available. For instance, although *P. putida* KT2440 is unable to use ethylene glycol as a sole carbon source, the genes encoding its degradation are present in KT2440, and multiple *P. putida* strains grow on this compound (92). Therefore, the incongruences still remaining in the model pave the way towards targeted identification of new genes responsible for orphan reactions, the deciphering of unknown regulatory mechanisms as well as a guide for future adaptive laboratory evolution (ALE) experiments.

#### Growth experiments on carbon and nitrogen sources

Individual colonies of *P. putida* KT2440 and mutant strains available at Pseudomonas Reference Culture Collection (PRCC) (60) were picked from the surface of cultures freshly grown on LB medium plates supplemented with 30 μg/ml of chloramphenicol, streaked onto M8 pre-growth medium plates (0.1% [wt/vol] glucose, 0.1 g/liter NH_4_Cl, 1 mM MgSO_4_, 0.6 mg/L Fe-citrate, and micronutrients), and grown overnight at 30°C. Pre-growth of cells on M8 pre-growth medium was sufficient to deplete nutrient reserves such that the subsequent growth assays with different carbon, nitrogen, and sulfur sources were dependent on the nutritional sources provided. The biomass of the overnight plates described above was recovered from the plate surface and suspended in 15 ml of M9 or M8 liquid medium (Daniels et al, 2010) to a turbidity at 660 nm (OD660) of 0.1. The wells of the microplates were filled with 180 μl of the cellular suspension, and 20 μl of each carbon, nitrogen, or sulfur source was added to reach a final concentration of 5 mM. For sulfur source assays, the MgSO_4_ in the medium was replaced with MgCl_2_. Positive-control wells consisted of full minimal medium containing glucose, NH_4_Cl, and MgSO_4_ as carbon, nitrogen, and sulfur sources, respectively; negative-control wells contained medium without cell inoculate.

All data recordings were performed using a type FP-1100-C Bioscreen C MBR analyzer system (OY Growth Curves Ab Ltd., Raisio, Finland) at 30°C, with continuous agitation. The turbidity was measured using a wideband filter at 420 to 580 nm every 60 min over a 24-h period. Each strain was assayed at least three times for each of the compounds tested, and plates were visually examined following each assay in order to verify the results.

#### Gene essentiality predictions on *i*LB and glucose

*In silico* LB medium (*i*LB) was formulate based on the composition of commercial LB medium and the conditional essential gene analysis in *P. putida* (61) (SI1). For predicting gene essentiality in glucose, a glucose minimal medium was simulated as described in SI1. The *singleGeneDeletion* function in the Cobra Toolbox (89) with minimization of metabolic adjustment (MOMA) algorithm (93) were used to simulate knockouts. A gene was considered to be essential if its removal reduced the growth rate below 10% of the growth rate in the original model. The gene essentiality analysis under environmental perturbations was performed analogously. Glucose, ammonium, inorganic phosphate, sulfate and Fe^2+^ were used as default carbon, nitrogen, phosphate, sulfur and iron sources, respectively. Growth in the *i*LB medium and nutrients used by *i*JN1411 (Figure 2) was simulated and the *singleGeneDeletion* function was used to identify the essential genes in each condition. For the stressor analysis, default glucose minimal medium was used and 34 chemical stressors including heavy metals e.g., Hg, Pb, SbO_3_, Cu, CrO_4_; organic solvents e.g., xylene, toluene; antibiotic e.g., tetracycline, chloramphenicol, ampicillin; ROS e.g., H_2_O_2_; and miscellanea compounds such as TNT and formaldehyde were introduced to the model using sink reactions (10 mmol/gDW·h) (Table S6).

#### Identification of robustness cycles

For the identification of metabolic cycles consuming ATP we applied an optimization algorithm developed by Pinchuk and collegues to enumerate ATP-dependent futile cycles (75). Briefly, an artificial ATP synthesis reaction (ADP + Pi + H^+^ → ATP + H2O) with positive flux is added to the model and all exchange fluxes are constrained to zero so that no metabolites can enter or exit the system. This approach ensures that the futile cycle(s) must take on non-zero fluxes in order to hydrolyze the ATP that is produced by the artificial ATP producing reaction. Futile cycles based on transport reactions and coupled kinases/phosphatases were subsequently omitted from the analysis.

### *P. putida* multi-strain genome-scale modeling

We constructed a gene orthology matrix between KT2440 and the ten sequenced *P. putida* strains (Table S5). In addition, we included *Pseudomonas entomophila* L48 in the analysis, a phylogenetically related organism, in order to extend the approach beyond of *P. putida*. We then identified the genes present in *i*JN1411 for which no orthologous gene was found in each of the strains analyzed, and subsequently removed the corresponding GPR from *i*JN1411 to obtain the strain-specific GENREs. This process was performed automatically following established procedures (17, 94).

## Acknowledgements

This work was supported by the Spanish Spanish Ministry of Economy and Competitiveness through the RobDcode grant (BIO2014-59528-JIN). Work in Abengoa Research was supported by grants from CDTI and The Empowerputida European Commission grant. The funders had no role in study design, data collection and interpretation, or the decision to submit the work for publication.

The authors thanks to J Monk his valuable help with the multiple correspondence analyses of available GENREs, to M. Abrams his critical reading of the manuscript, R. van Heck for providing *i*EB1050 model, and B. Jiménez for helping in Figure 3.

## Funding Information

MINECO: Spanish Ministry of Economy and Competitiveness. BIO2014-59528-JIN Juan Nogales

EU: European Union’s Horizon 2020: No 635536

Juan L. Ramos

## Author Contribution

JN conceived and designed the study, carried out the reconstructions, performed the constraint-based analyses, analyzed the data and drafted the manuscript. BOP contributed to the design the study, analyzing the data and writing the manuscript. SG contributed to the computational analysis. JLR and ED performed the growth experiments and knockout analyses and analyzed the data. All the authors contributed to the final version of the manuscript.

## Supplementary Material

SI1: Supplementary Information, including additional text, figures and references (PDF)

Table S1: Model and Manual Curation (.xlsx)

Table S2: Table S2_Nutrients and Carbon flux Validations (.xlsx)

Table S3: Biomass Formulation (.xlsx)

Table S4: Essentiality Analysis (.xlsx)

Table S5: Multi strains Modeling (.xlsx)

Table S6: Robustness Breakdown (.xlsx)

Table S7: Futile Cycles Identification (.xlsx)

Table S8: Gene Expression of Robustness Cycles (.xlsx)

